# Factors affecting CRISPR-Cas defense against antibiotic resistance plasmids harbored by *Enterococcus faecalis* laboratory model strains and clinical isolates

**DOI:** 10.1101/2025.03.10.642232

**Authors:** Tahira Amdid Ratna, Belle M. Sharon, Cesar Alejandro Barros Velin, Kelli L. Palmer

**Author notes:** **Corresponding author: Dr. Kelli Palmer:**.

## Abstract

*Enterococcus faecalis* is a Gram-positive bacterium and opportunistic pathogen that acquires resistance to a wide range of antibiotics by horizontal gene transfer (HGT). The rapid increase of multidrug-resistant (MDR) bacteria including MDR *E. faecalis* necessitates the development of alternative therapies and a deeper understanding of the factors that impact HGT. CRISPR-Cas systems provide sequence-specific defense against HGT. From previous studies, we know that *E. faecalis* CRISPR-Cas provides sequence-specific anti-plasmid defense during agar plate biofilm mating and in the murine intestine. Those studies were mainly conducted using laboratory model strains with a single, CRISPR-targeted plasmid in the donor. MDR *E. faecalis* typically possess multiple plasmids that are diverse in sequence and may interact with each other to impact plasmid transfer and CRISPR-Cas efficacy. Here, we altered multiple parameters of our standard *in vitro* conjugation assays to assess CRISPR-Cas efficacy, including the number and genotype of plasmids in the donor; and laboratory model strains as donor versus recent human isolates as donor during conjugation. We found that the plasmids pTEF2 and pCF10, which are not targeted by CRISPR-Cas in our recipient, enhance the conjugative transfer of the CRISPR-targeted plasmid pTEF1 into both wild-type and CRISPR-Cas-deficient (via deletion of *cas9*) recipient cells. However, the effect of pTEF2 on pTEF1 transfer is much more pronounced, with a striking 6-log increase in pTEF1 conjugation frequency when pTEF2 is also present in the donor and recipients are deficient for CRISPR-Cas (compared to 4-log for pCF10). Overall, this study provides insight about the interplay between plasmids and CRISPR-Cas defense, opening avenues for developing novel therapeutic strategies to curb HGT among bacterial pathogens, and highlighting pTEF2 as a plasmid for additional mechanistic study.

**IMPORTANCE:** The emergence of MDR bacteria, including MDR *E. faecalis,* limits treatment options and necessitates development of alternative therapeutics. In these circumstances, bacterial CRISPR-Cas systems are being explored by the field to develop CRISPR-based antimicrobials. However, in many cases CRISPR-Cas efficacy has only been assessed using laboratory model strains. More studies are required that investigate clinical isolates, as those are the intended targets for CRISPR antimicrobials. Here, we demonstrated how the number of plasmids harbored by an *E. faecalis* donor strain can affect the apparent efficacy of CRISPR-Cas anti-plasmid defense in a recipient strain. Overall, our research is important to develop improved CRISPR-based antimicrobials to combat the spread and accumulation of antibiotic resistance determinants.

## INTRODUCTION

*E. faecalis* is a Gram-positive opportunistic pathogen (1, 2). Despite being a natural inhabitant of the mammalian gastrointestinal tract, due to fecal contamination, this pathogen is frequently found in soil, sewage, water, and food (3, 4). *E. faecalis* is also a leading cause of hospital-acquired infections in the United States, especially in immunocompromised patients (5). *E. faecalis* is considered a serious threat by the United States Centers for Disease Control and Prevention due to high occurrence of resistance to a variety of antibiotics including vancomycin, a last resort antibiotic, leaving few treatment options (6–8).

Horizontal gene transfer (HGT) disseminates antibiotic resistance genes among bacterial pathogens, including *E. faecalis* (9). Many studies have identified mobile genetic elements (MGEs) such as pheromone-responsive plasmids (PRPs), mobilizable plasmids, and transposons as means of HGT in *E. faecalis* (9–11). Conjugation is the most studied form of HGT in *E. faecalis* (12–15). PRPs are conjugative plasmids that can achieve very high conjugation frequencies due to their transfer mechanism which capitalizes on sex pheromone production by recipient cells to facilitate cell contact with donors (16–18). Thus, *E. faecalis* donor strains harboring a PRP can transfer the plasmid to a recipient cell via conjugation and produce a transconjugant (16, 19). PRPs were first identified in *E. faecalis* and appear to exclusively replicate within the enterococci (20, 21). Examples of well-studied, model PRPs are pAD1 and pCF10 (20, 22, 23).

Clustered regularly interspaced short palindromic repeat and CRISPR associated proteins (CRISPR-Cas) systems, can provide sequence-specific genome defense against HGT (24–26). This system preserves a genetic memory of past encounters with MGEs via short sequences called spacers. Upon spacer-dependent recognition of a targeted MGE, Cas-encoded nucleases can cleave and deactivate it (26–28). In *E. faecalis*, predominantly type IIA CRISPR-Cas systems are found and characterized by the presence of type II specific nuclease Cas9 (formerly known as Csn1) (29–31). Genome analysis has also identified presence of type IIC CRISPR-Cas system in *E. faecalis* recently (32). Thus far, four different CRISPR-Cas loci namely CRISPR1-Cas, CRISPR2, CRISPR3-Cas and CRISPR4 have been identified in *E. faecalis* (29, 30, 33). The plasmid recipient strain used in this study, *E. faecalis* urinary tract isolate T11RF (34, 35) possesses a native CRISPR3-Cas system (30). The T11RF CRISPR3-Cas spacer 6 has sequence identity to the model PRP pAD1 and reduces acquisition frequency of this plasmid in T11RF (30, 34).

From bioinformatic studies, we know that many MDR *E. faecalis* are “immunocompromised” and lack functional CRISPR-Cas systems, which likely allows for accumulation of antibiotic resistance-encoding MGEs in these strains (30, 36, 37). Using the non-MDR *E. faecalis* T11RF as a model recipient, previous works demonstrated that *E. faecalis* CRISPR-Cas confers defense against pAD1 derivatives both *in vitro* and *in vivo* (in the murine intestine) (34, 38). These experimental studies established conclusively that CRISPR-Cas can serve as an anti-plasmid defense system in *E. faecalis*. Yet, a limitation of these studies is that they assessed CRISPR-Cas efficacy against model antibiotic resistance plasmids and used laboratory model donor strains derived from the natively plasmid-free 1975 oral isolate OG1 (39). *E. faecalis* MDR clinical isolates can possess up to six plasmids (2, 40, 41), most of which have not been investigated beyond sequencing. These *E. faecalis* clinical isolates are plasmid reservoirs and donors from which antibiotic resistance spreads. Theoretically, plasmids in these donor cells may collaborate to enhance their transfer rates and/or overcome anti-plasmid defense in the recipient cell. Conjugation efficiency is influenced by a complex interplay of intracellular and intercellular interactions between distinct plasmids (42). Indeed, it has also been reported that anti-CRISPR phages can cooperate to defeat CRISPR-Cas defense in host bacteria (43). Hence, the number and genotype of plasmids harbored by an *E. faecalis* donor are factors that need to be explored to determine their impact on CRISPR-Cas anti-plasmid defense. Other recent research has also demonstrated that the effectiveness of a CRISPR-based antimicrobial against *E. faecalis* fecal surveillance isolates can be affected by competitive factors (such as bacteriocins) produced by the isolates (44). This emphasizes the need to incorporate clinical isolates in research to understand CRISPR-Cas efficacy against them.

Overall, in this study we incorporated both laboratory model strains and clinical isolates to examine the correlation between the number of plasmids present in a donor cell and the ability of CRISPR-Cas to enact its defense mechanism in the recipient. We found that presence of multiple plasmids in a donor strain and the lack of active CRISPR-Cas system in a recipient strain can additively confer strikingly high conjugative transfer frequencies of antibiotic resistance plasmids. Our results can be leveraged to develop enhanced CRISPR-based antimicrobials. Our research also underscores the importance of investigating additional potential factors that may influence the effectiveness of CRISPR-Cas anti-plasmid defense.

## RESULTS

### CRISPR-Cas defense in the recipient is less effective when pTEF1 and pTEF2 are present in the donor strain, compared to pTEF1 alone

We first examined the correlation between the number of plasmids present in the donor cell and the ability of CRISPR-Cas in the recipient cell to enact its defense mechanism. We hypothesized that the presence of multiple plasmids in an *E. faecalis* donor would increase the conjugative transfer of antibiotic resistance and negatively impact CRISPR-Cas genome defense in recipients. To generate isogenic plasmid donors with varying number of plasmids, we used *E. faecalis* clinical isolate V583 as the plasmid donor and *E. faecalis* laboratory model strain OG1SSp (16, 45) as the recipient strain in a conjugation assay. We chose V583 because it was among the first vancomycin-resistant clinical isolates reported in the United States (46) and contains 3 plasmids, pTEF1, pTEF2 and pTEF3 (2). pTEF1 is a PRP encoding erythromycin resistance (2). pTEF2 is also a PRP but does not encode antibiotic resistance genes (2). pTEF1 and pTEF2 share sequence features with the model PRPs pAD1 (47) and pCF10 (18), respectively, which respond to different peptide pheromones (2). pTEF3 is a nonconjugative broad host range plasmid (2). Conjugation assays were conducted as described previously (34) (Figure 1). Antibiotic-containing agar plates were used to select for OG1SSp(pTEF1) transconjugants. For all the conjugation assays, conjugation frequency was calculated by dividing the CFU/mL of the transconjugants by the CFU/mL of the donors. Eight transconjugants were selected for PCR to confirm their plasmid content. Two of these transconjugants had one plasmid - OG1SSp(pTEF1) - and 6 transconjugants had two plasmids - OG1SSp(pTEF1, pTEF2). Transfer of pTEF3 was not observed (Figure S1). Thus, we generated our first controlled set of donors with varying number of plasmids.

**Figure 1:**
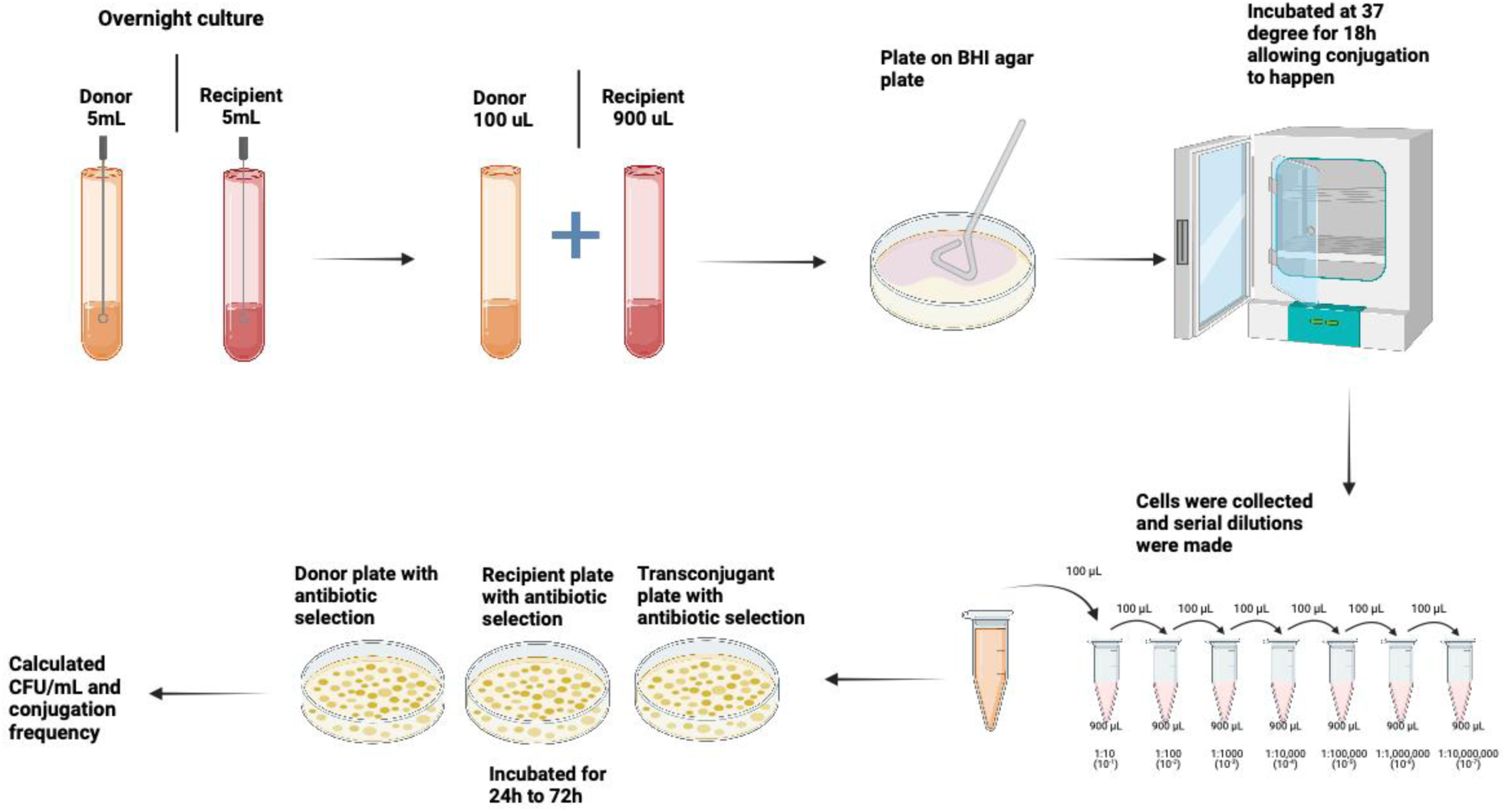
Overview of conjugation assays. In day 1, donor and recipients were inoculated in 5 mL BHI without antibiotic selection and incubated overnight at 37°C. The next day, the cultures were diluted 1:10 into fresh BHI and incubated for 1.5 h at 37°C or until log phase reached. A 100 μL volume of donor culture was mixed with 900 μL volume of recipient culture, and the mixture was pelleted at 13,000 rpm for 1 min. 900 μL supernatant was discarded, and the remaining 100 μL of supernatant was used to resuspend the pellet. This was then plated on a BHI agar plate and incubated at 37°C for 18 h. Cells were collected from the plate with 2 mL 1X PBS supplemented with 2 mM EDTA. This was serially diluted (10^-1^ to 10^-8^). Dilutions were plated on BHI agar plates supplemented with appropriate antibiotics to quantify donors, recipients and transconjugants. CFU/mL was calculated after 48 h to 72 h incubation at 37°C and used for calculating conjugation frequency (34) (Created in https://BioRender.com).

We used the newly established OG1SSp(pTEF1) and OG1SSp(pTEF1, pTEF2) donors in further conjugation assays (Figure 2A). *E. faecalis* urinary tract isolate T11RF (34, 35) was used as the recipient strain for these experiments (Figure 2A). T11RF is vancomycin-susceptible and often used in comparative analyses with V583 (11, 30, 34). Though T11RF and V583 share 99.5% nucleotide identity in their core genome, T11RF lacks ∼620 kb of horizontally acquired genome content present in V583 (11, 30). Spacer 6 of T11RF CRISPR3-Cas system has sequence identity with a region in pTEF1; none of the spacers have sequence identity with pTEF2, indicating that this plasmid is not targeted by the T11RF CRISPR3-Cas system. The other two recipients for this study were T11RFΔ*cas9,* where the *cas9* gene was deleted to deactivate the CRISPR-Cas system, and T11RFΔ*cas9*+CR3, where the *cas9* gene was complemented back to the Δ*cas9* mutant into a neutral site on the chromosome (34) (Figure 2A). Conjugation assays were conducted as described previously (34) (Figure 1). As expected, T11RF CRISPR3-Cas provided genome defense against pTEF1 for both OG1SSp(pTEF1) and OG1SSp(pTEF1,pTEF2) donors, as significantly higher pTEF1 conjugation frequency was observed in the absence of *cas9* for both (Figure 2B-a,2B-b).

**Figure 2:**
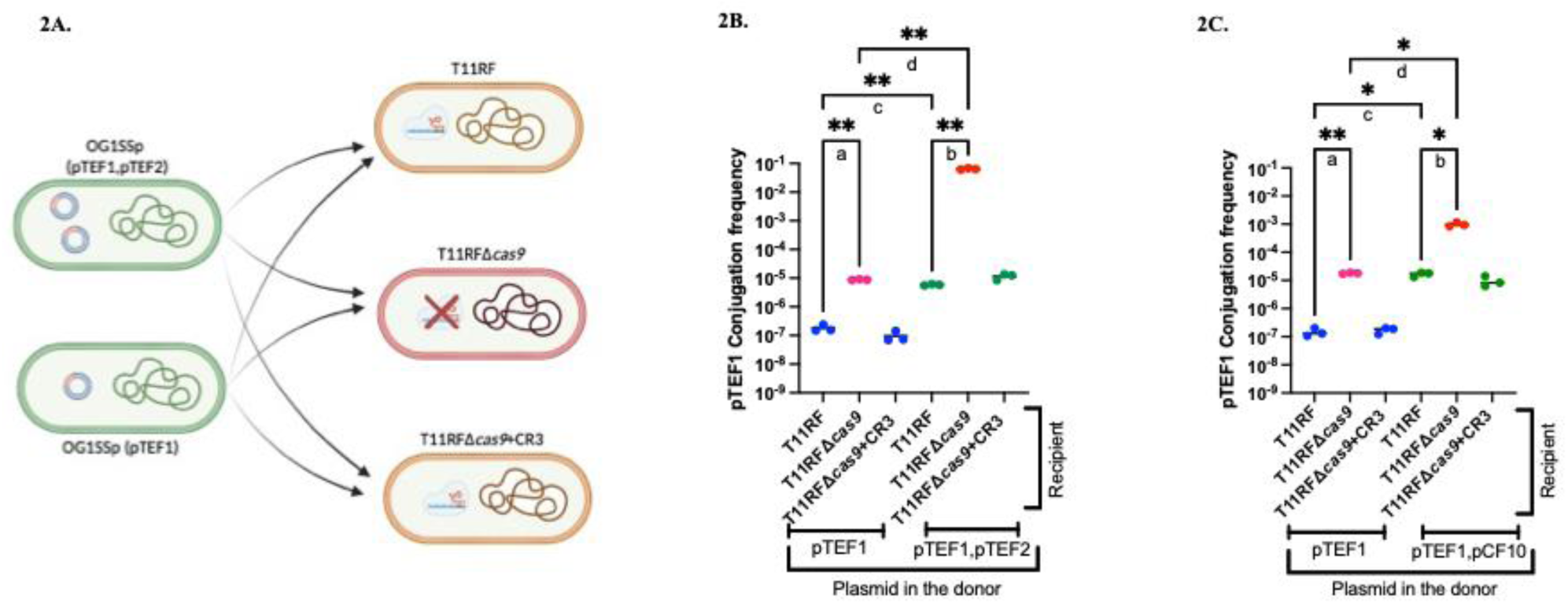
T11RF CRISPR3-Cas efficacy against pTEF1 for OG1SSp(pTEF1) and OG1SSp(pTEF1,pTEF2) or OG1SSp(pTEF1,pCF10) donor strains. Figure 2A shows the donors OG1SSp(pTEF1, pTEF2) or OG1SSp(pTEF1) and the recipients T11RF, T11RFΔ*cas9*, and T11RFΔ*cas9*+CR3 that are used in the conjugation assays for figure 2B (Created in https://BioRender.com). In Figure 2B and 2C, conjugation frequency of pTEF1 and statistical analysis are showing. T11RF CRISPR3-Cas can provide sequence specific genome defense against OG1SSp (pTEF1) and OG1SSp (pTEF1,pTEF2) (2B-a, 2B-b) (Dunnett’s T3 multiple comparisons test, \*\**P*-value = 0.0029 (2B-a), \*\**P*-value = 0.0067 (2B-b)). However, higher conjugation frequency of pTEF1 was observed when the donor strain had both pTEF1 and pTEF2 present, compared with only pTEF1 (2B-c, 2B-d) (Dunnett’s T3 multiple comparisons test, \*\**P*-value = 0.0049 (2B-c), \*\**P*-value = 0.0067 (2B-d)). T11RF CRISPR3-Cas can also provide sequence specific genome defense against OG1SSp (pTEF1) and OG1SSp (pTEF1,pCF10) (2C-a, 2C-b) (Dunnett’s T3 multiple comparisons test, \*\**P*-value = 0.0053 (2C-a), \**P*-value = 0.0360 (2C-b)). Higher conjugation frequency of pTEF1 was observed when the donor strain had both pTEF1 and pCF10 present, compared with only pTEF1 (2C-c, 2C-d) (Dunnett’s T3 multiple comparisons test, \**P*-value = 0.0438 (2C-c), \**P*-value = 0.0361 (2C-d)).

We then compared the pTEF1 conjugation frequency for the two donors. We expected that, if CRISPR-Cas was equally effective against pTEF1 irrespective of pTEF2 presence, the pTEF1 conjugation frequency would be unchanged for wild-type T11RF recipients. However, we observed a 32-fold increase in pTEF1 conjugation frequency for wild-type T11RF when the donor strain had both pTEF1 and pTEF2 present, compared with only pTEF1 (Figure 2B-c). Moreover, in the case of T11RFΔ*cas9* as the recipient strain, a striking 7278-fold increase in pTEF1 conjugation frequency was observed when the donor strain had both pTEF1 and pTEF2 present, compared with only pTEF1 (Figure 2B-d). We analyzed the plasmid content of 6 T11RF and 6 T11RFΔ*cas9* transconjugants using PCR, finding that both pTEF1 and pTEF2 transferred to all 6 T11RF transconjugants and to 5 of 6 T11RFΔ*cas9* transconjugants (Figure S2).

Our results support our hypothesis and demonstrate that CRISPR-Cas efficacy against pTEF1 is diminished when another plasmid, pTEF2, is present in the donor cell and co-transfers with pTEF1. Moreover, pTEF1 transfer frequency is extraordinarily high (a 6-log increase) when donors possess both pTEF1 and pTEF2, and recipients lack functional CRISPR-Cas defense compared to donor where only pTEF1 is present and the recipient possesses active CRISPR-Cas.

### pCF10 increased pTEF1 conjugation frequency when both plasmids were present together in a donor

pCF10 is a well-studied model PRP that shares identical conjugation genes with pTEF2 (2, 18). pCF10 encodes tetracycline resistance and is not targeted by T11RF CRISPR3-Cas (34). Therefore, we explored whether pCF10 can “help” pTEF1 in a manner similar to that observed for pTEF2. We generated another donor strain, OG1SSp(pTEF1, pCF10), and then used OG1SSp(pTEF1) and OG1SSp(pTEF1, pCF10) donor strains in conjugation assays. As expected, T11RF CRISPR3-Cas provided genome defense against pTEF1 from both donors, as higher pTEF1 conjugation frequencies were observed in the absence of *cas9* (Figure 2C-a, 2C-b).

We then compared the pTEF1 conjugation frequency for the two donors. A 114-fold increase in pTEF1 conjugation frequency was observed when the donor strain had both pTEF1 and pCF10 present, compared with only pTEF1 (Figure 2C-c). In the case of T11RFΔ*cas9* as the recipient strain, a 54-fold increase in pTEF1 conjugation frequency was observed when the donor strain had both pTEF1 and pCF10 present, compared with only pTEF1 (Figure 2C-d). During the conjugation assay, we only tracked pTEF1 using erythromycin-containing agar. Hence, we screened transconjugants for the presence of both pTEF1 and pCF10 plasmids using erythromycin- and tetracycline-containing agars, respectively. For all transconjugants screened, pTEF1 and pCF10 plasmids were transferred together (Figure S3). Overall, from these observations we can say that pCF10 also increased pTEF1 conjugation frequency when present together in a donor.

We noted that Figure 2B (for effect of pTEF2 in donors) and 2C (for effect of pCF10 in donors) were nearly superimposable, with the exception of the results for T11RFΔ*cas9* recipients. The presence of pTEF2 in donors conferred an additional 2-log increase in pTEF1 conjugation frequency into T11RFΔ*cas9* recipients, compared to the presence of pCF10 in donors (Figure 2B-2C). pTEF2 possesses 21 genes (in two clusters) that pCF10 lacks (Supplemental Dataset S1). Conserved domain analysis identified putative functions including plasmid partitioning, pheromone response, and modification-dependent DNA nicking for 15 of the genes; no conserved domains were identified for 6 of the genes. Further experimentation will be required to establish the basis for the extraordinarily high pTEF2-assisted pTEF1 transfer rate for T11RFΔ*cas9* recipients.

### Deletion of aggregation substance gene *prgB* from pCF10 decreases pTEF1 conjugation frequency from a multi-plasmid donor

When we observed that pCF10 was able to increase pTEF1 conjugation frequency when present together in a donor, we wanted to explore which region of pCF10 might be important for this plasmid cooperativity. The pCF10 gene *prgB* encodes the aggregation substance protein Asc10 and plays an important role during conjugative transfer of pCF10 via intracellular aggregation or clumping (16, 48, 49). From previous studies, we know that deletion of *prgB* contributes to decreased virulence and also decreased conjugation frequency of pCF10 in *E. faecalis* (48, 50, 51). Hence, we obtained a previously reported variant of pCF10 with the aggregation substance gene *prgB* deleted, referred to as pCF10-8 (48). We generated the donor strain OG1SSp(pTEF1, pCF10-8) and then used OG1SSp(pTEF1) and OG1SSp(pTEF1, pCF10-8) donor strains in conjugation assays. CRISPR-Cas provided genome defense against pTEF1 from both donors, as higher pTEF1 conjugation frequency was observed in the absence of *cas9* in recipient (Figure 3A-a, 3A-b). When we compared the conjugation frequency of pTEF1 from the two donors, we still observed a 21-fold increase in conjugation frequency of pTEF1 to recipient T11RF (Figure 3A-c) or a 7-fold increase to T11RFΔ*cas9* (Figure 3A-d), when the donor strain had both pTEF1 and pCF10-8 present, compared with only pTEF1. This means that even with the deletion of aggregation substance gene *prgB*, pCF10-8 can still increase pTEF1 conjugation frequency. However, for wild type pCF10 we observed higher fold increase in conjugation frequency of pTEF1 (Figure 2C-c, 2C-d) than pCF10-8. Thus, we concluded that the pCF10 aggregation substance gene *prgB* is important for the increased conjugation frequency of pTEF1 from a multi-plasmid donor, but it is not the only important region under the conditions tested.

**Figure 3:**
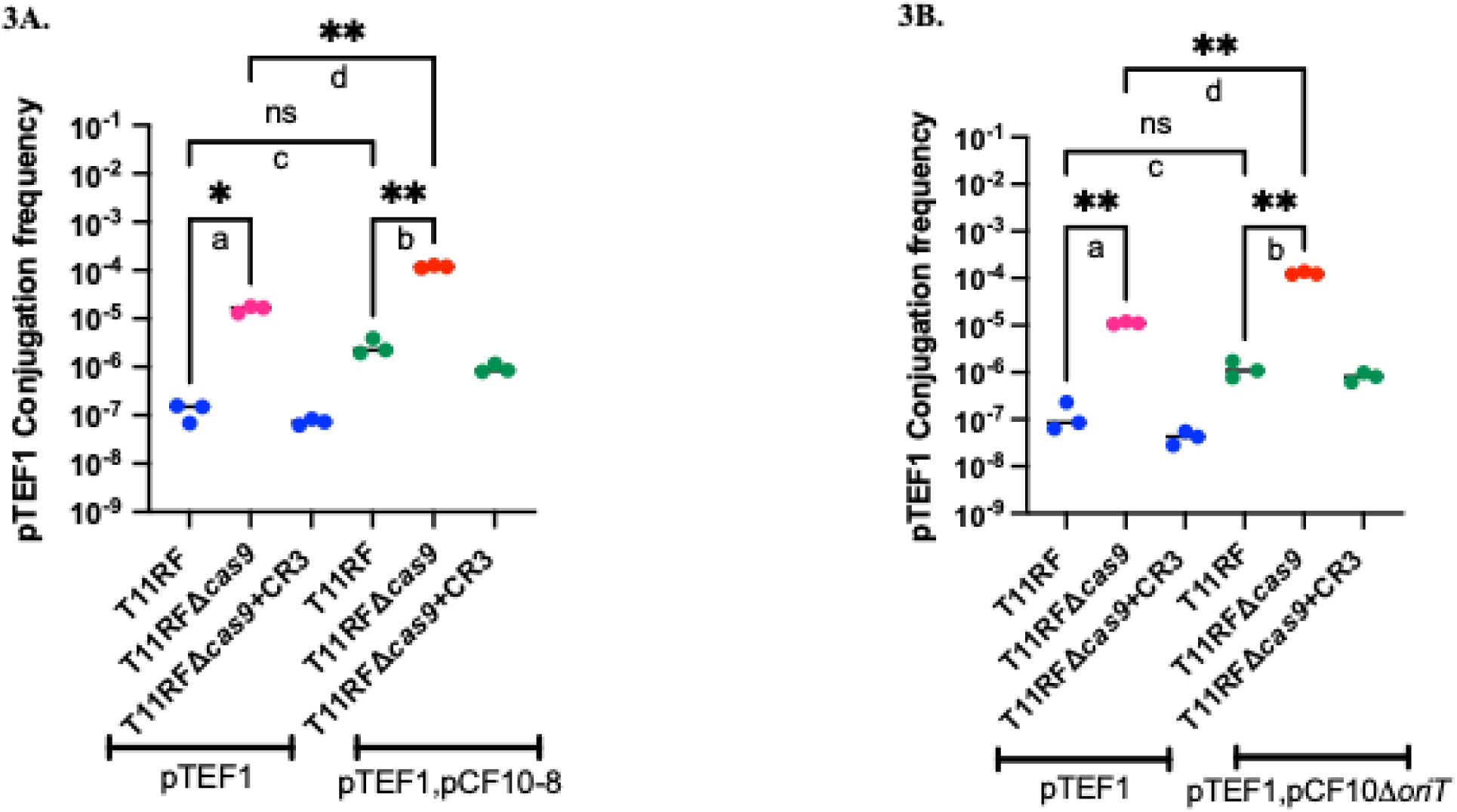
T11RF CRISPR3-Cas efficacy against pTEF1 plasmid in OG1SSp(pTEF1) and OG1SSp(pTEF1,pCF10-8) or OG1SSp(pTEF1,pCF10Δ*oriT*) donor strains. Figure 3A and 3B show the conjugation frequency of pTEF1 and statistical analysis. CRISPR-Cas was able to provide sequence specific genome defense against pTEF1 in both donors as higher conjugation frequency was observed in the absence of *cas9* in recipients (3A-a, 3A-b) (Dunnett’s T3 multiple comparisons test, \**P*-value = 0.0358 (3A-a), \*\**P*-value = 0.0064 (3A-b)). Higher conjugation frequency was observed when the donor strain had both pTEF1 and pCF10-8 present, compared with only pTEF1 to T11RF (3A-c) or T11RFΔ*cas9* (3A-d) (Dunnett’s T3 multiple comparisons test, ns *P*-value = 0.2139 (3A-c), \*\**P*-value = 0.0088 (3A-d)). Even with the deletion of aggregation substance gene *prgB*, pCF10 plasmid can still increase the conjugation frequency of pTEF1. CRISPR-Cas was able to provide sequence specific genome defense against pTEF1 in both donors as higher conjugation frequency was observed in the absence of *cas9* in recipients (3B-a, 3B-b) (Dunnett’s T3 multiple comparisons test, \*\**P*-value = 0.0063 (3B-a), \*\**P*-value = 0.0074 (3B-b). Higher conjugation frequency was observed when the donor strain had both pTEF1 and pCF10Δ*oriT* present, compared with only pTEF1 to T11RF (3B-c) or T11RFΔ*cas9* (3B-d) (Dunnett’s T3 multiple comparisons test, ns *P*-value = 0.2598 (3B-c), \*\**P*-value = 0.0087 (3B-d)). Even with the deletion *oriT*, pCF10 plasmid can still increase pTEF1 conjugation frequency.

### Deletion of *oriT* from pCF10 decreases pTEF1 conjugation frequency from a multi-plasmid donor

The origin of transfer region, or *oriT,* is the region on plasmid DNA where conjugation initiates, hence playing an important role during conjugative transfer of plasmids (52). A previous study demonstrated that deletion of *oriT* from pCF10 significantly decreases its conjugation frequency (53). Therefore, we obtained an *oriT*-deleted mutant of pCF10, pCF10Δ*oriT* (53), and generated the donor strain OG1SSp (pTEF1, pCF10Δ*oriT*). We used OG1SSp (pTEF1) and OG1SSp (pTEF1, pCF10Δ*oriT*) donor strains in conjugation assays with the same recipients as before. CRISPR-Cas provided genome defense against pTEF1 from both donors, as higher pTEF1 conjugation frequency was observed in the absence of *cas9* in recipients (Figure 3B-a, 3B-b). When we compared the conjugation frequency of pTEF1 in two donors we still observed a 9-fold increase in conjugation frequency of pTEF1 to recipient T11RF (Figure 3B-c) or a 11-fold increase to T11RFΔ*cas9* (Figure 3B-d), when the donor strain had both pTEF1 and pCF10Δ*oriT* present, compared with only pTEF1. These results were similar to what we observed with pCF10-8 and indicate that even with the deletion *oriT*, pCF10Δ*oriT* can still increase pTEF1 conjugation frequency. However, for wild type pCF10 we observed higher fold increase in conjugation frequency of pTEF1 (Figure 2C-c, 2C-d) than pCF10Δ*oriT*. Thus, we concluded that, the *oriT* of pCF10 is also important for increased conjugation frequency of pTEF1 from a multi-plasmid donor. The potential additive effect of *prgB/oriT* double deletion was not investigated in our study but would be informative.

### T11RF CRISPR3-Cas can target clinical isolates and provide defense against resistance plasmids harbored by those clinical isolates

We included 10 previously reported hospital fecal surveillance isolates (44) and 25 previously reported urine isolates (41) in this study. At first, we checked if spacers in the T11RF CRISPR3-Cas system could target the chromosomes or plasmids of these clinical isolates. We found that 6 of 10 fecal surveillance isolates (Supplemental Dataset S2) and 17 of 25 urine isolates (Supplemental Dataset S2) were targeted by the T11RF CRISPR3-Cas system as evidenced by 100% nucleotide sequence identity between spacer and target, and the presence of a suitable protospacer adjacent motif (previously defined in reference 34). The targeted plasmids and antibiotic resistance genes encoded by those plasmids are shown in Table 1 and Supplemental Dataset S2. Moving forward, we will denote plasmids targeted by the T11RF CRISPR3-Cas system in bold text (for each isolate, we denote their plasmid content by their sizes in kilobases [kb] after the isolate name).

**Table 1:** Fecal surveillance isolates and urine isolates for conjugation assays. Five fecal surveillance isolates (the first five listed) and three urine isolates (the last three) used for conjugation assays in this study are presented in Table 1. Targeted plasmids or chromosome are shown in bold in column 1, T11RF CRISPR3-array spacer that target the plasmid, or the chromosome are shown in column 2 and resistance gene encoded in the targeted plasmids or chromosome are shown in bold in column 3. The rep family of the plasmids are shown in column 4 and the CRISPR-targeted plasmid rep family is indicated in bold (41, 44). Isolates for which conjugation was not detected are highlighted in grey.

| Fecal surveillance isolates and urine isolates | T11RF CRISPR3-array spacer that targets the plasmid or chromosome of the isolates | Resistance gene encoded in the targeted plasmids or chromosome | Rep family of the plasmids |
| --- | --- | --- | --- |
| 59 (88kb, **43kb**) | Spacer 6 | *vanHAX*, *ermB* | rep9c, **rep9a** |
| 43-2 (**54kb**) | Spacer 6 | *ermB* | **rep9a**, rep2 |
| 2-1 (**149kb**) | Spacer 6 | *ermB* | **rep9a** |
| 133-1 (**172kb**, 43kb) | Spacer 7 | *ermB* | Unknown, Unknown |
| 101-1 (**65kb**) | Spacer 7 | *ermB* | **rep9b** |
| EfsC8 (**52kb**) | spacer 7 | *ermB* | **rep9c** |
| EfsC33 (**104kb**, 72kb) | spacer 6 | *ermB* | **rep9a**, rep9b |
| EfsC61 (**57kb**) | spacer 7 | *ant(6)-la* | **rep2** |

The fecal isolates 59 (88 kb**, 43 kb**), 43-2 (**54 kb**), 2-1 (**149 kb**), 133-1 (**172 kb**, 43 kb), 101-1 (**65 kb**) and urine isolates EfsC8 (**52 kb**), EfsC33 (**104 kb**, 72 kb), EfsC61 (**57 kb**) were used as plasmid donors in conjugation assays with T11RF recipients (Table 1). We chose these clinical isolates as plasmid donors because they harbor CRISPR3-targeted resistance plasmids, and thus we were able to track those plasmids during conjugation assays. Plasmids with sequences targeted by spacer 6 and spacer 7 are shown in Figure 4 and Figure 5, respectively. For fecal isolates 59 (88kb, **43kb**), 43-2 (**54kb**), 101-1 (**65Kb**) and urine isolate EfsC61 (**57kb**), conjugation was not detected. Fecal isolates 2-1 (**149 kb**), 133-1 (**172kb**, 43kb) and urine isolates EfsC8 (**52kb**), EfsC33 (**104 kb,** 73kb) conjugated with our recipients at detectable levels. The T11RF CRISPR3-Cas system provided defense against the **149 kb** plasmid from the fecal isolate 2-1 (Figure 4A) with a more modest effectiveness against the **104kb** plasmid from the urine isolate EfsC33 (Figure 4B). Interestingly, this was very comparable to T11RF CRISPR3-Cas spacer 7 targeted plasmids (Figure 5). The T11RF CRISPR3-Cas system provided defense against the **172 kb** plasmid in fecal isolate 133-1(Figure 5A) and a weaker protective effect against the **52 kb** plasmid in urine isolate EfsC8 (Figure 5B) as higher conjugation frequencies for these plasmids were observed in the absence of *cas9* in recipients.

**Figure 4:**
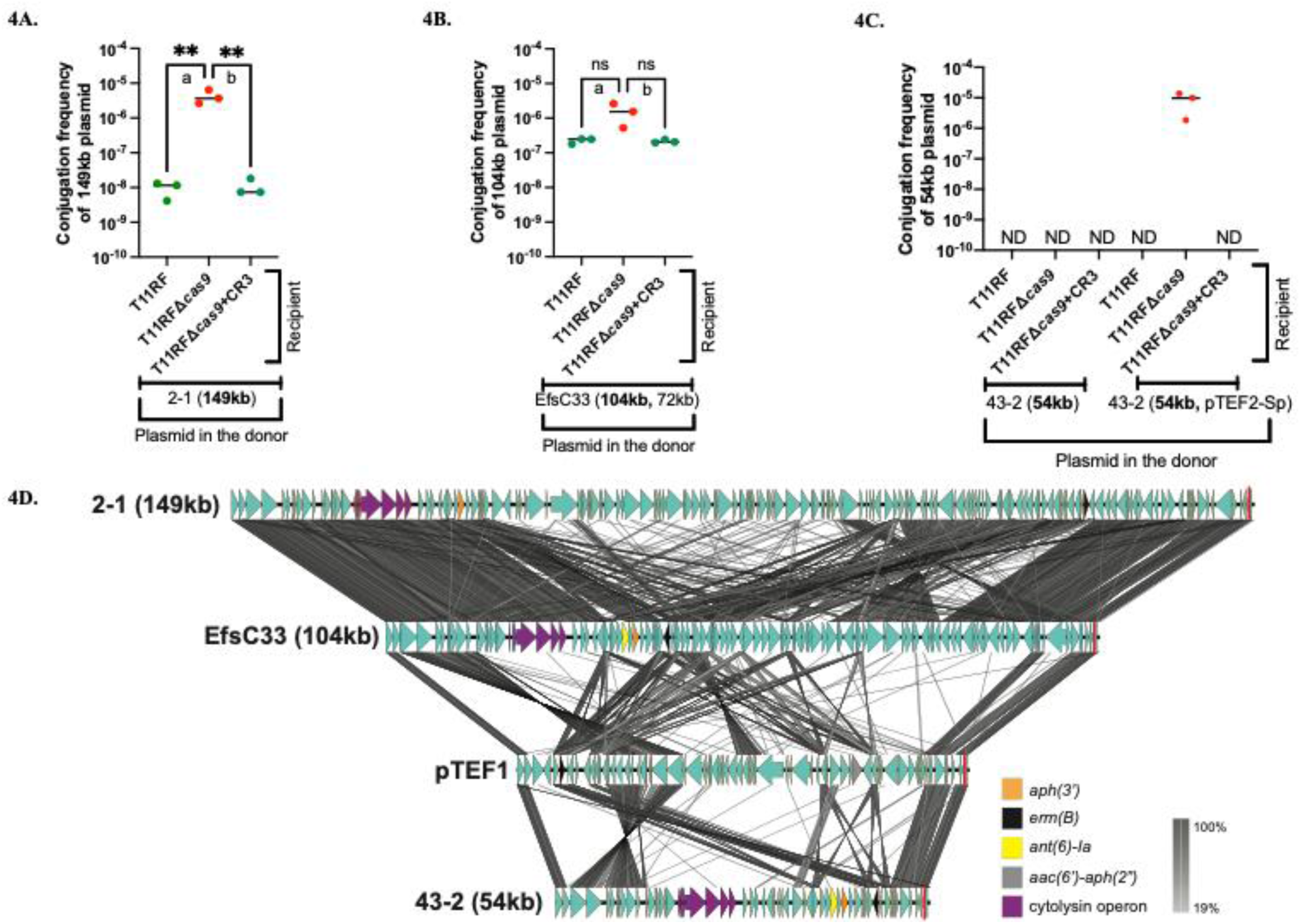
Clinical isolate plasmid donors targeted by T11RF CRISPR3-Cas spacer 6. T11RF CRISPR3-Cas decreased the rate of transfer of the **149 kb** resistance plasmid from fecal surveillance isolate 2-1 (4A) (Tukey’s multiple comparisons test, \*\**P*-value = 0.0088 (4A-a,4A-b)) and the **104 kb** plasmid from urine isolate EfsC33 (4B) (Tukey’s multiple comparisons test, ns *P*-value = 0.0786 (4B-a), ns *P*-value = 0.0765 (4B-b)). Conjugation was not detected for **54 kb** plasmid from fecal isolate 43-2 with the recipients when present alone in the donor. pTEF2-Sp mobilized the **54kb** plasmid from fecal isolate 43-2 in the absence of *cas9* in the recipient (4C). tblastx alignment of plasmids targeted by T11RF CRISPR3 spacer 6 sequence is presented in 4D. Location of spacer 6 sequence is indicated with a red line. Coding sequences are denoted by arrows with antibiotic resistance genes and cytolysin operon genes indicated in unique colors. Lines drawn between plasmid sequences show sequence identity (%) (4D).

**Figure 5:**
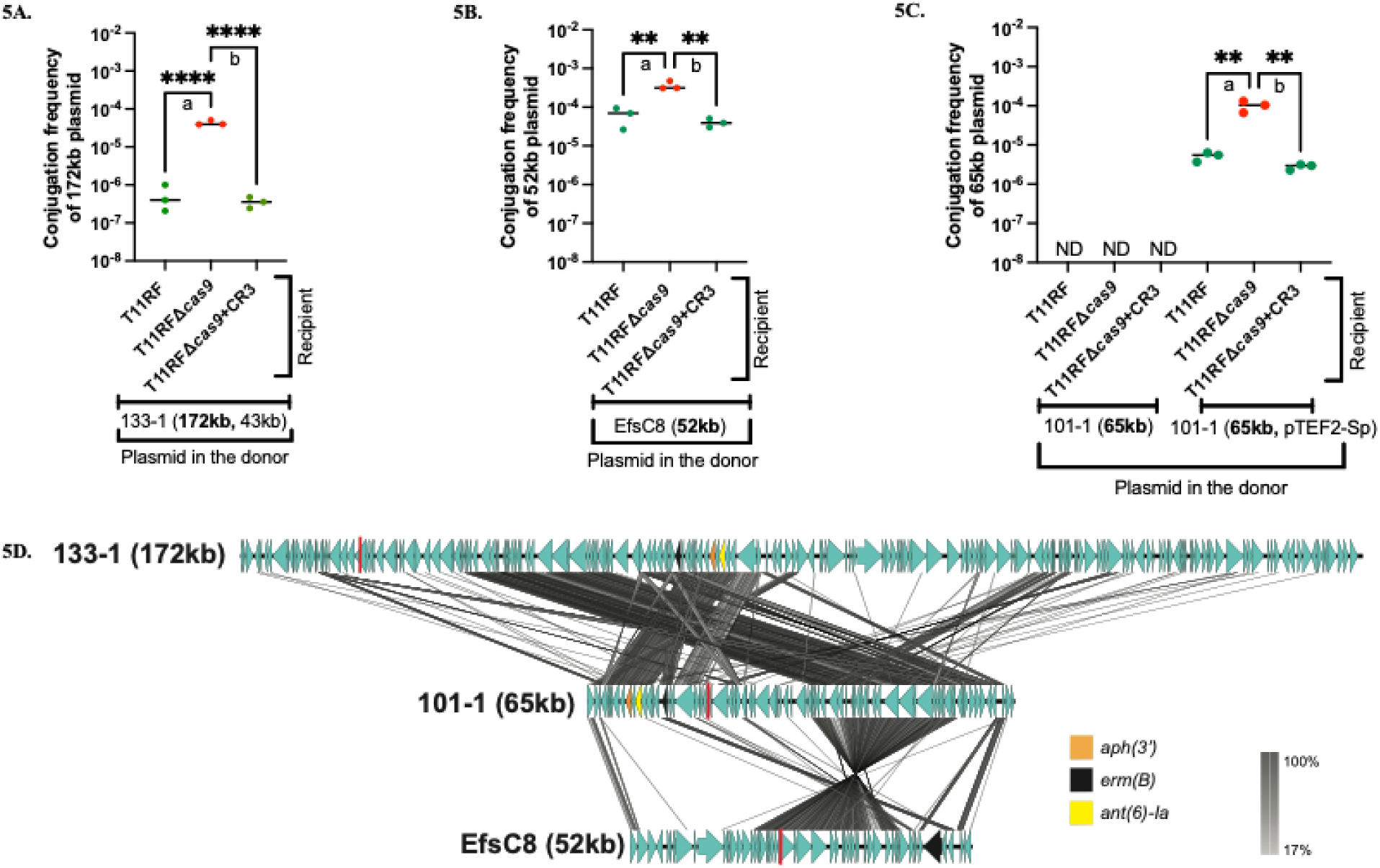
Clinical isolate plasmid donors targeted by T11RF CRISPR3-Cas spacer 7. T11RF CRISPR3-Cas was able to decrease the rate of transfer of the resistance plasmid **172kb** from clinical fecal surveillance isolate 133-1 (5A) (Tukey’s multiple comparisons test, \*\*\*\**P*-value < 0.0001 (5A-a,5A-b)) and **52kb** from clinical urine isolate EfsC8 (5B) (Tukey’s multiple comparisons test, \*\**P*-value = 0.0016 (5B-a), \*\**P*-value = 0.0011 (5B-b)). Conjugation was not detected for fecal isolate 101-1 with the recipients when present alone in the donor. pTEF2-Sp plasmid mobilized the clinical resistant plasmid **65kb** from the fecal isolate 101-1. However, T11RF CRISPR3-Cas was still able to decrease the rate of transfer of the resistance plasmid **65kb** as higher conjugation frequency was observed in the absence of *cas9* gene (5C) (Tukey’s multiple comparisons test, \*\**P*-value = 0.0019 (5C-a), \*\**P*-value = 0.0017 (5C-b)). tblastx alignment of plasmids targeted by T11RF CRISPR3 spacer 7 sequence is presented in 5D. Location of spacer 7 sequence is indicated with a red line. Coding sequences are denoted by arrows with antibiotic resistance genes indicated in unique colors. Lines drawn between plasmid sequences show sequence identity (%).

We noted highly variable transfer rates of the CRISPR3-targeted plasmids and the magnitude of CRISPR3-Cas efficacy against them, despite the plasmids being PRPs from similar plasmid rep families. We performed plasmid alignment of the spacer 6- and spacer 7-targeted plasmids (Figures 4D and 5D, respectively). Our alignments corroborate conclusions reached by prior investigation of the *E. faecalis* urine isolate collection, namely, that plasmid rep typing fails to capture the genetic diversity of PRPs (41). The variability of CRISPR3-Cas defense against these “wild” plasmids is likely due to their genetic diversity and interactions with other MGEs in the donor, which together influence plasmid transfer frequency and interaction with recipient defense systems. This study marks a significant milestone as the first to demonstrate the efficacy of the T11RF CRISPR3-Cas system in targeting plasmids from recent clinical isolates.

### pTEF2 increases conjugation frequency of resistance plasmids from two fecal isolates and helps them escape T11RF CRISPR3-Cas defense

Conjugation was not detected for the fecal isolates 43-2 (**54kb**), and 101-1 (**65kb**) with T11RF (Figure 4C, Figure 5C). We investigated what effect pTEF2 would have on the transfer of these plasmids, given our results demonstrating that pTEF2 could assist pTEF1 with transfer and apparent evasion of CRISPR-Cas defense (Figure 2B). We made new donor strains with pTEF2-Sp (pTEF2 modified to encode spectinomycin resistance (54)). We used these new donor strains, 43-2 (**54kb**, pTEF2-Sp), and 101-1 (**65kb**, pTEF2-Sp) in conjugation assays with our previous recipients. We observed that pTEF2-Sp can mobilize the clinical resistant plasmids, as they conjugated to the recipients. However, T11RF CRISPR3-Cas still provided defense against the **54 kb** plasmid of the 43-2 strain (Figure 4C) and the **65 kb** plasmid of the 101-1 strain (Figure 5C), as in the absence of T11RF *cas9,* higher conjugation frequencies were observed. For the 43-2 donor strain, conjugation of the **54 kb** plasmid was detected only when *cas9* was deleted from the recipient strain. We did not perform the same experiment with pCF10, as pCF10 encodes tetracycline resistance, and 43-2 and 101-1 also encode tetracycline resistance in their chromosome. Therefore, we concluded that pTEF2 can increase conjugation frequency of other resistance plasmids and help them in escaping T11RF CRISPR3-Cas defense. Our findings underscore the importance of considering plasmid cooperativity when refining CRISPR-based antimicrobial designs. Our work suggests that targeted depletion of pTEF2 from *E. faecalis* populations may specifically be useful for mitigating antibiotic resistance plasmid spread.

## DISCUSSION

In this era of antibiotic resistance, *E. faecalis* is considered a serious threat as it can acquire a large number of resistance genes via HGT (2, 9). Bacterial CRISPR-Cas systems may be leveraged as an alternative treatment approach, providing sequence-specific cleavage of antibiotic resistance plasmids in *E. faecalis* (34, 38). Most available studies of CRISPR-Cas efficacy have used laboratory model strains and laboratory model plasmids. However, clinical isolates can behave differently than the laboratory model strains, and CRISPR-Cas efficacy can be affected by many factors (44). Therefore, for this study we included both laboratory model strains and clinical isolates to study previously unexplored factors that might affect CRISPR-Cas efficacy. Our hypothesis was that the number of plasmids present in a donor strain is an important factor that might affect CRISPR-Cas efficacy. Through our experiments we showed our hypothesis is supported and found two plasmid pairs (i) pTEF1,pTEF2 and (ii) pTEF1, pCF10 that showed plasmid cooperativity. From our experiments we saw pTEF2 or pCF10 were able to increase pTEF1 conjugation frequency. We identified two regions, (i) aggregation substance gene *prgB* and (ii) *oriT* of pCF10, potentially important for the plasmid cooperativity as deletion of these regions resulted in decreased conjugation frequency of pTEF1.

We included 10 clinical fecal surveillance isolates (44) and 25 urine isolates (41) in this study. We found that 6 fecal surveillance isolates, and 17 urine isolates can be targeted by the T11RF CRISPR3-Cas system. When we used these clinical isolates as plasmid donors, we found that T11RF CRISPR3-Cas was able to provide defense against the resistance plasmids harbored by those isolates. To the best of our knowledge, this is the first research studying T11RF CRISPR3-Cas efficacy against a collection of clinical isolates. Interestingly, there is a significant impact of CRISPR-Cas on targeted plasmids, but the magnitude of the impact differed for different clinical isolates. There were a few clinical isolates that did not conjugate with our recipient at detectable levels. However, when we conjugated pTEF2-Sp plasmid in those clinical isolates, pTEF2-Sp was able to mobilize two clinical resistant plasmids (i) 54kb, pTEF2-Sp from 43-2 and (ii) 65kb, pTEF2-Sp from 101-1. However, T11RF CRISPR3-Cas system was still able to provide defense against those plasmids once they were mobilized. This finding is similar to our plasmid cooperativity results of pTEF1, pTEF2 and pTEF1, pCF10. Together these plasmid cooperativity results indicate that the first conjugative resistance plasmid lacks something essential for robust transfer, to which they are getting “help” from the second plasmid, and together, they escape T11RF CRISPR3-Cas defense by dint of higher conjugation frequency. A limitation of our experimental design is, we did not pinpoint the exact mechanism of this plasmid cooperativity. It could be the helper plasmid enhancing the transfer frequency of the first plasmid, the helper plasmid inhibiting the CRISPR-Cas defense against the first plasmid, or both.

Overall, we conclude that the presence of multiple interacting plasmids in donor strains and the lack of an active CRISPR-Cas system in a recipient strain act additively to confer extremely high plasmid transfer frequencies in *E. faecalis*. To date, a few multi-plasmid studies have demonstrated how ecological settings influence the maintenance of individual plasmids within a multi-plasmid community, how a conjugative helper plasmid can mobilize a non-conjugative virulence plasmid, and how interactions between plasmids can affect conjugation efficiency (42, 55, 56). Hence, our study opens a new direction of plasmid cooperativity and contributes towards designing novel CRISPR antimicrobials.

Apart from the plasmid number present in donor strains, there may be other factors affecting CRISPR-Cas efficacy, such as biofilm formation capacity or internal CRISPR-Cas regulation in the recipient strains. For example, transcriptional regulation of *cas9* was reported as a major factor which can influence CRISPR-Cas genome defense in *S. pyogenes* (57). Hence, it is necessary to continue to study potential factors that might affect CRISPR-Cas efficacy, in future studies. Overall, our research is significant in its application towards designing improved CRISPR-based antimicrobials as an alternative approach to combat HGT and antibiotic resistance. Our research also emphasizes to study factors affecting CRISPR-Cas efficacy and conducting research using clinical isolates. The more information we gather on the efficacy of CRISPR-Cas, the more effectively we can harness it as an alternative treatment approach.

## MATERIALS AND METHODS

### Bacterial strains and reagents used

Strains and plasmids used in this study are shown in Table S1. *E. faecalis* strains were cultured in brain heart infusion (BHI) broth or agar at 37°C without shaking, unless otherwise stated. Antibiotics were added at the following concentrations: rifampin (R), 50 μg/mL; fusidic acid (F), 25 μg/mL; streptomycin (S), 500 μg/mL; tetracycline (Tet), 10 μg/mL; spectinomycin (Sp), 500 μg/mL; erythromycin (Erm), 50 μg/mL; vancomycin (Van), 10 μg/mL; gentamicin (Gent), 10 μg/mL. Routine PCR analysis was performed using *Taq* polymerase (New England Biolabs). Primers (Sigma-Aldrich) used in this study are shown in Table S2.

### Conjugation assays

Conjugation assays were conducted as previously described (34). CFU/mL was determined using the following formula: CFU/mL = number of colony/(amount plated (mL) x dilution factor). Conjugation frequency was calculated by dividing the CFU/mL of the transconjugants by the CFU/mL of the donors. Raw CFU/mL data for conjugation experiments are shown in Supplemental Dataset S3. Graphs were prepared and statistical analysis performed using GraphPad Prism (Version 9.0.0).

### Selection of fecal surveillance isolates and urine isolates

10 fecal surveillance isolates (44), and 25 urine isolates (41) were analyzed for this study. First, a database was created in Geneious with the previously reported genome sequences of the 10 fecal surveillance isolates or 25 urine isolates. All CRISPR3 spacer sequences from T11RF were queried against the database using BLASTn. 6 of 10 fecal surveillance isolates and 17 of 25 urine isolates were identified with T11RF CRISPR3-Cas targets in either plasmids or in the chromosome with 100% nucleotide sequence identity (Supplemental Dataset S2). The presence of the expected protospacer adjacent motif sequence (34) adjacent to the target sequence was also confirmed.

### Generation of 43-2 (54kb, pTEF2-Sp) and 101-1 (65kb, pTEF2-Sp) donor strains and conjugation assays

*E. faecalis* V19 (pTEF2-Sp) (54) was used as donor and fecal isolates 43-2 (**54kb**), 101-1 (**65kb**) were used as recipients in conjugation assays to generate 43-2 (**54kb**, pTEF2-Sp) and 101-1 (**65kb**, pTEF2-Sp) donor strains. PCR was performed to confirm the plasmids present in the transconjugant. Once confirmed, 43-2 (**54kb**, pTEF2-Sp) and 101-1 (**65kb**, pTEF2-Sp), were used as donors in conjugation assays using T11RF, T11RFΔ*cas9* and T11RFΔ*cas9*+CR3 as recipients. Conjugation assays were done in biological triplicate using the previously described method (34). Antibiotic selection plates were prepared according to the resistance gene present in the donor and recipients (Table S1). All the CFU/mL calculation and data analysis were done as described above.

### Plasmid alignment and analysis

Plasmid tblastx alignments were performed using EasyFig 3.0.0 at default parameters. Antimicrobial resistance genes were identified using ABRicate v1.0.1 querying the ResFinder database at default parameters. pTEF2-specific genes (in comparison to pCF10) were analyzed using NCBI Conserved Domains (58), PSORTb 3.0.3 (59), and InterPro (60). Plasmid replicon types were predicted using PlasmidFinder v2.1 at default parameters. Where plasmid replicon type was not identified using PlasmidFinder, plasmids were analyzed using NCBI Conserved Domains to identify the *rep* gene. Further typing was completed using blastn and PlasmidFinder queries at 70% identity threshold (41).

## ACKNOWLEDGMENTS

We thank Dr. Julia Willett for providing us pCF10Δ*oriT* and pCF10-8 and Dr. Breck Duerkop for providing us pTEF2-Sp. We also thank Dr. Nicole J. De Nisco for providing us the clinical urine isolates.

## SUPPLEMENTAL MATERIALS

**Figure S1:**
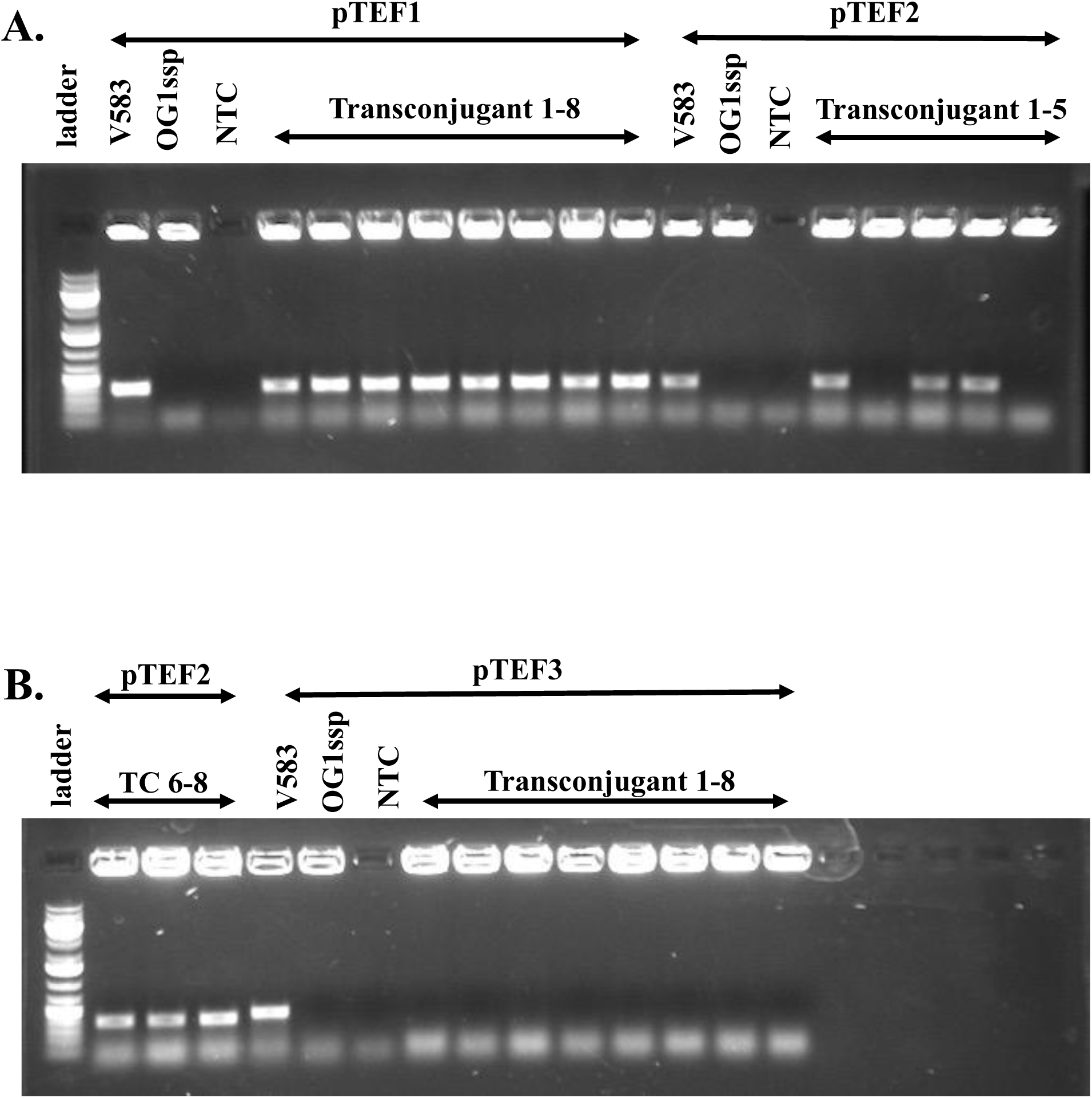
Identification of plasmid pTEF1, pTEF2 and pTEF3 by PCR. Primers designed for pTEF1, pTEF2 and pTEF3 were used to screen the transconjugants. V583 was used as the positive control, and the amplified plasmid is indicated in each case. The gel image shows that it contains all three plasmids. OG1SSp was used as the negative control and the amplified plasmid is indicated in each case. The gel image shows that it contains none of the plasmid. A no template control (NTC) was also added in each case. The gel image shows that all eight selected transconjugants contain the pTEF1 plasmid (A); six transconjugants contain the pTEF2 plasmid (A,B) and none of the transconjugants contain pTEF3 plasmid (B).

**Figure S2:**
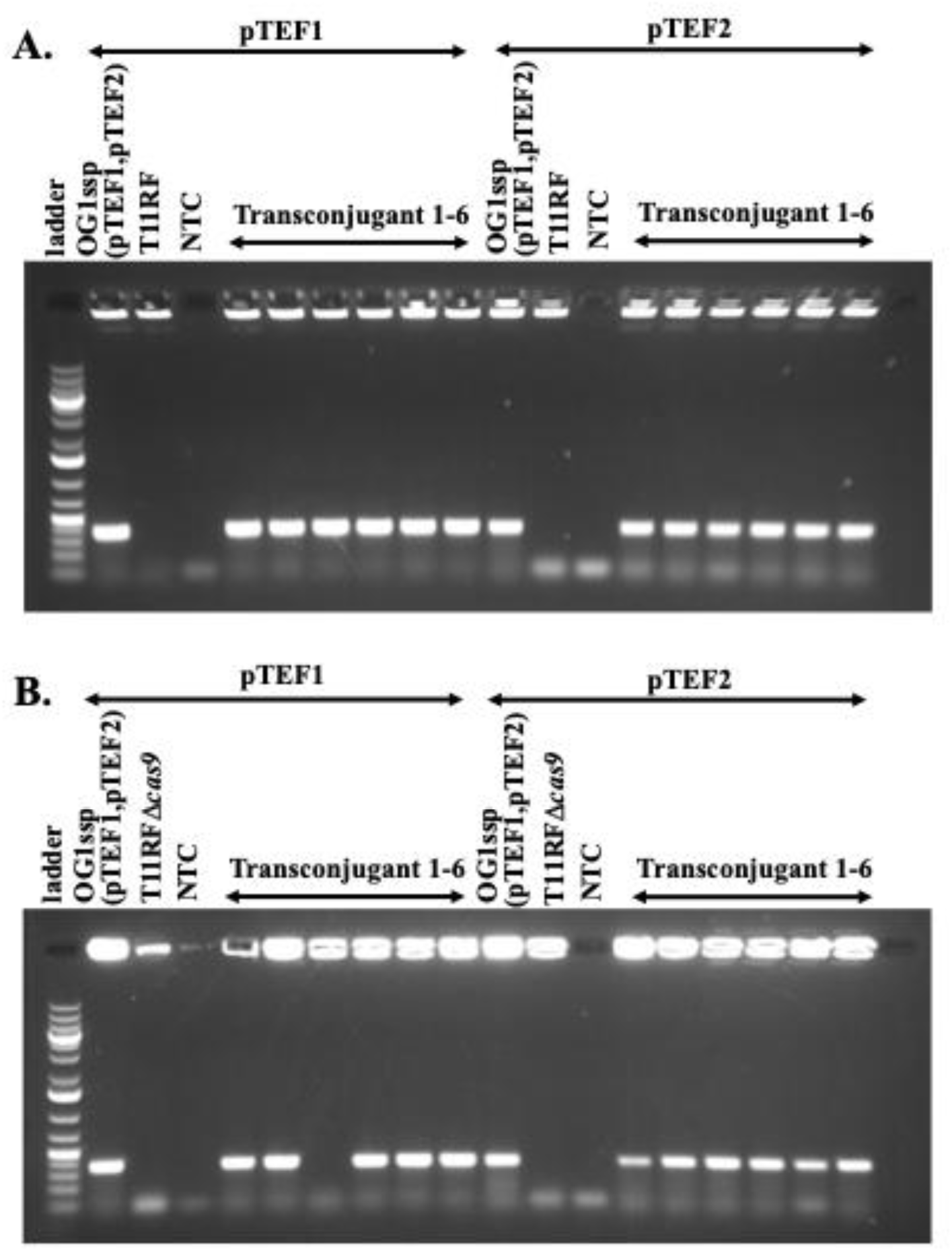
Identification of plasmid pTEF1 and pTEF2 in transconjugants by PCR. Primers designed for pTEF1 and pTEF2 were used to screen the transconjugants. OG1SSp (pTEF1, pTEF2) was used as the positive control, and the amplified plasmid is indicated in each case. The gel image shows that it contains both pTEF1 and pTEF2 plasmids (A, B). T11RF (A) and T11RFΔ*cas9* (B) were used as the negative control and gel image shows that they contain none of the plasmids. A no template control (NTC) was also added in each case. The gel image shows that all the six selected transconjugants contain pTEF1 and pTEF2 plasmids when T11RF was the recipient (A). Five transconjugants contain pTEF1 plasmid and all six transconjugants contain pTEF2 plasmid when T11RFΔ*cas9* was the recipient (B).

**Figure S3:**
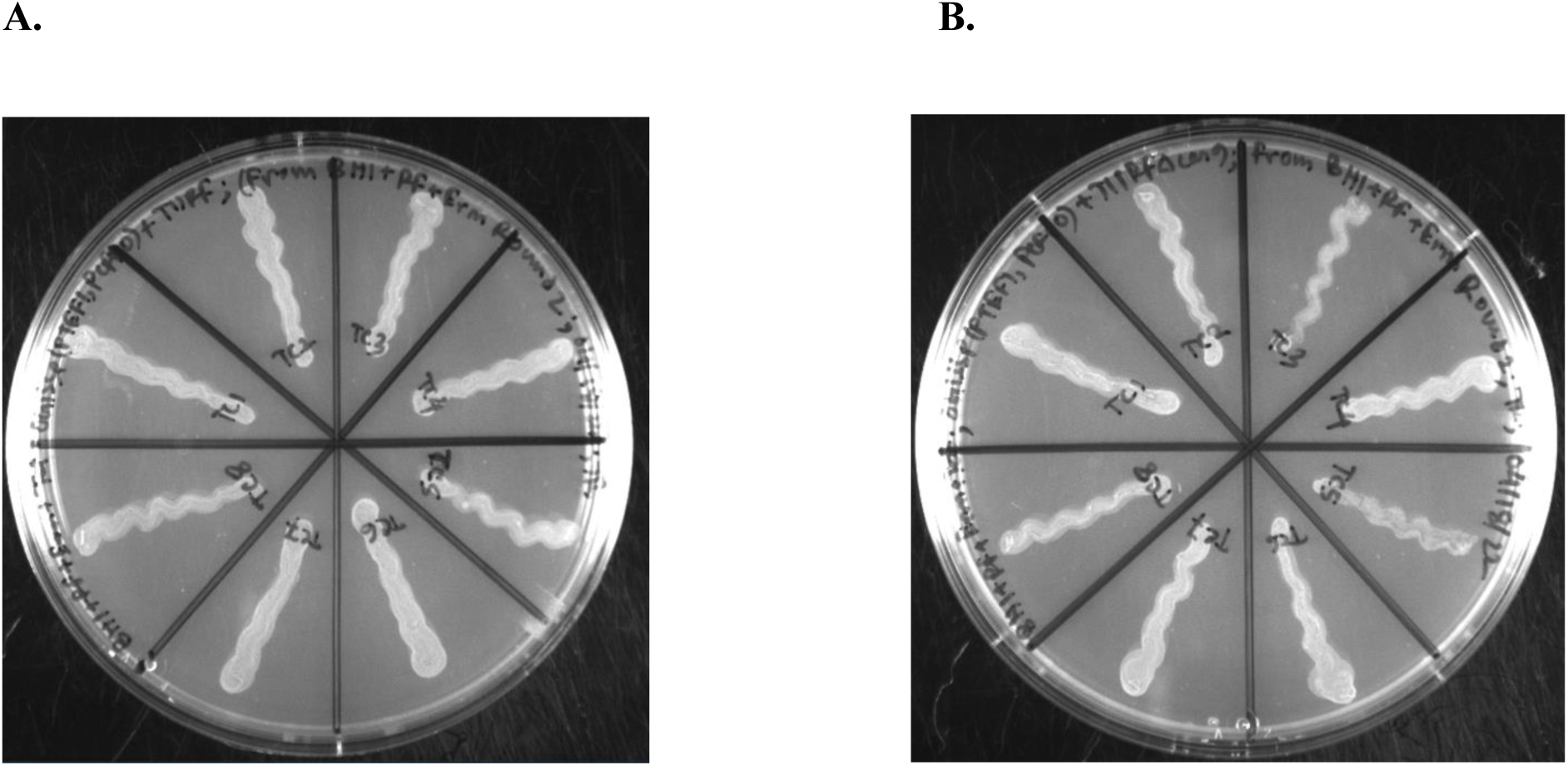
Identification of plasmid pTEF1 and pCF10 in transconjugants. 8 transconjugants were picked from the original transconjugant plates and restreaked where OG1SSp (pTEF1,pCF10) was donor and either T11RF (A) or T11RFΔ*cas9* (B) were recipients. In the original transconjugant plate only pTEF1 was selected by erythromycin antibiotic. In the restreaked plate both pTEF1 and pCF10 were selected by erythromycin and tetracycline respectively. We found that in all cases pTEF1 and pCF10 were transferred together.

**Table S1:** *E. faecalis* strains and plasmids used in this study.

| Strain | Description | Reference(s) |
| --- | --- | --- |
| V583 | Vancomycin-resistant *E. faecalis* clinical isolate harboring pTEF1, pTEF2 and pTEF3 plasmids | (2, 46) |
| OG1SSp | Spectinomycin- and streptomycin-resistant derivative of OG1 *E. faecalis* strain harboring no plasmids | (16, 45) |
| OG1SSp (pTEF1) | Spectinomycin- and streptomycin-resistant derivative of OG1 *E. faecalis* strain harboring erythromycin resistant pTEF1 plasmid | This study |
| OG1SSp (pTEF1,pTEF2) | Spectinomycin- and streptomycin-resistant derivative of OG1 *E. faecalis* strain harboring pTEF1 and pTEF2 plasmids | This study |
| T11RF | Rifampin- and fusidic acid-resistant derivative of *E. faecalis* urine isolate T11 | (34, 35) |
| T11RFΔcas9 | T11RF CRISPR3-cas9 deletion mutant | (34) |
| T11RFΔcas9+CR3 | T11RFΔcas9 mutant with chromosomal integration of CRISPR3 cas9 between EFMG_00904 and EFMG_00905 | (34) |
| OG1RF (pCF10) | Rifampin- and fusidic acid-resistant derivative of OG1 *E. faecalis* strain harboring pCF10 plasmid encoding tetracycline resistance on Tn925 | (16, 61, 62) |
| OG1SSp (pTEF1,pCF10) | Spectinomycin- and streptomycin-resistant derivative of OG1 *E. faecalis* strain harboring pTEF1 and pCF10 | This study |
| CK104 (pCF10-8) | OG1RFΔupp strain harboring pCF10-8 plasmid (*prgB* deletion mutant retaining first three and last three codons, Asc10-) | (48, 63) |
| OG1SSp (pTEF1,pCF10-8) | Spectinomycin- and streptomycin-resistant derivative of OG1 *E. faecalis* strain harboring pTEF1 and pCF10-8 plasmids | This study |
| CK104 (pCF10ΔoriT) | OG1RFΔupp strain harboring pCF10ΔoriT plasmid. | (53) |
| OG1SSp (pTEF1, pCF10ΔoriT) | Spectinomycin- and streptomycin-resistant derivative of OG1 *E. faecalis* strain harboring pTEF1 and pCF10ΔoriT | This study |
| 59 (88kb,43kb) | Fecal surveillance *E. faecalis* isolate 59 harboring vancomycin resistant 88 kb plasmid and erythromycin resistant 43 kb plasmid | (44) |
| 43-2 (54kb) | Fecal surveillance *E. faecalis* isolate 43-2 harboring erythromycin resistant 54kb plasmid | (44) |
| 2-1 (149kb) | Fecal surveillance *E. faecalis* isolate 2-1 harboring erythromycin resistant 149kb plasmid | (44) |
| 133-1 (172kb) | Fecal surveillance *E. faecalis* isolate 133-1 harboring erythromycin resistant 172kb plasmid | (44) |
| 101-1 (65kb) | Fecal surveillance *E. faecalis* isolate 101-1 harboring erythromycin resistant 65kb plasmid | (44) |
| 142-1 (77kb, 59kb) | Fecal surveillance *E. faecalis* isolate 142-1 harboring erythromycin resistant 77kb plasmid and 59kb plasmid | (44) |
| V19 (pTEF2-Sp) | Plasmid-free derivative of V583 Em<sup>S</sup>, Gm<sup>S</sup> harboring Spectinomycin resistant pTEF2 plasmid | (54) |
| 43-2 (54kb, pTEF2-Sp) | Fecal surveillance *E. faecalis* isolate 43-2 harboring erythromycin resistant 54kb plasmid and Spectinomycin resistant pTEF2 plasmid | This study |
| 101-1 (65kb, pTEF2-Sp) | Fecal surveillance *E. faecalis* isolate 101-1 harboring erythromycin resistant 65kb plasmid Spectinomycin resistant pTEF2 plasmid | This study |
| EfsC8 (52kb) | *E. faecalis* urine isolate EfsC8 harboring erythromycin resistant 52kb plasmid | (41) |
| EfsC33 (104kb, 72kb) | *E. faecalis* urine isolate EfsC33 harboring erythromycin resistant 104kb plasmid and 72kb plasmid | (41) |
| EfsC61 (57kb) | *E. faecalis* urine isolate EfsC61 harboring streptomycin resistant 57kb plasmid | (41) |

**Table S2:**
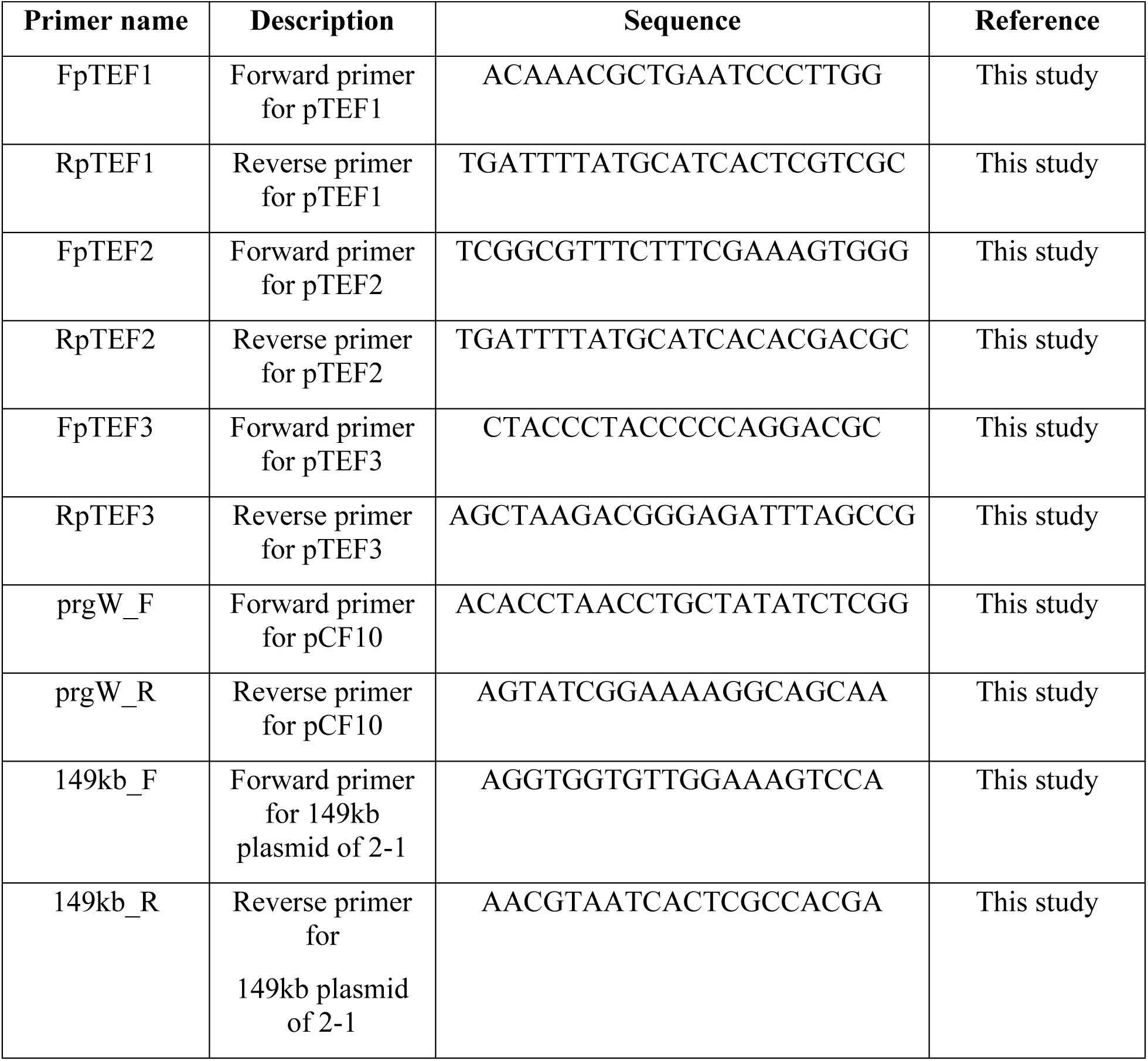
Primers used in this study.

